# Interpolation of Microbiome Composition in Longitudinal Datasets

**DOI:** 10.1101/2024.04.23.590847

**Authors:** Omri Peleg, Elhanan Borenstein

## Abstract

The human gut microbiome significantly impacts health, prompting a rise in longitudinal studies that capture microbiome samples at multiple time points. Such studies allow researchers to characterize microbiome changes over time, but importantly, also present major analytical challenges due to incomplete or irregular sampling. To address this challenge, longitudinal microbiome studies often employ various interpolation methods, aiming to infer missing microbiome data. However, to date, a comprehensive assessment of such microbiome interpolation techniques, as well as best practice guidelines for interpolating microbiome data, are still lacking.

This work aims to fill this gap, rigorously implementing and systematically evaluating a large array of interpolation methods, spanning several different categories, for longitudinal microbiome interpolation. To assess each method and its ability to accurately infer microbiome composition at missing time points, we used three longitudinal microbiome datasets that follow individuals over a long period of time, and a leave-one-out approach.

Overall, our analysis demonstrated that the K-nearest neighbors algorithm consistently outperforms other methods in interpolation accuracy, yet, accuracy varied widely across datasets, individuals, and time. Factors such as microbiome stability, sample size, and the time gap between interpolated and adjacent samples significantly influenced accuracy, allowing us to develop a model for predicting the expected interpolation accuracy at a missing time point.

Our findings, combined, suggest that accurate interpolation in longitudinal microbiome data is feasible, especially in dense cohorts. Furthermore, using our predictive model, future studies can interpolate data only in time points where the expected interpolation accuracy is high.

## Introduction

The human gut microbiome, the ensemble of microbial species that inhabit the human gut, undergoes substantial shifts in composition over time. These shifts are hard to predict and occur at different rates and in response to different perturbations. To better characterize such dynamics, recent years have seen an increase in the number of longitudinal studies of the human gut microbiome^1–6^. In these studies, individuals are sampled at multiple time points, in the hope of shedding light on temporal changes in the microbiome and on the correlation between compositional shifts and various health conditions or perturbations. Kostic *et al*.^5^, for example, have shown that the microbiome tends to undergo a significant change with the onset of diabetes in children. Similarly, several studies have shown a marked change in microbiome composition in response to dietary changes^7–9^, or during the first year of life^4,10^.

However, despite the importance of such studies, current longitudinal studies suffer from several drawbacks that make them challenging to analyze. First, most of these studies, unfortunately, include a relatively small number of individuals. Similarly, only a few studies follow individuals over a long period of time (*i*.*e*., more than a few weeks or months) or collect a large number of temporally dense samples from each participant. Moreover, since many bacteria are relatively rare and can be found only in a small number of samples, the resulting taxonomic profiles tend to be very sparse and challenging to analyze via standard statistical methods. Finally, in many cases samples are not being collected for all individuals at all time-points, or are being collected at different times. Such missing (or longitudinally mismatched) samples may be due to study design that is constrained by limited budget or personnel, partial compliance of participants, or technical issues. This, in turn can prevent the application of certain algorithms or statistical analyses, especially given the potentially limited number of participants and short study period mentioned above.

To address this latter challenge, microbiome studies have often resorted to interpolating the microbiome composition at missing time points, aiming to generate a more complete and uniform data for downstream analysis^4,11–13^. However, such an interpolation process has been carried out using a plethora of techniques, with most studies applying either spline interpolation^4,11,12^ or relatively simple and naïve approaches such as using the average abundance or the abundance at the last time point available in lieu of missing samples^13,14^. Furthermore, even though several more sophisticated interpolation techniques have been previously suggested^15,16^ (e.g., using temporal microbiome dynamics modeling methods to interpolate microbiome composition^17,18^), these techniques have been rarely used or assessed in practice. More importantly, while many possible approaches towards longitudinal microbiome interpolation can be considered, a comprehensive understanding of the optimal interpolation approach and of the factors that impact interpolation accuracy, as well as clear guidelines for interpolating samples in future studies, are currently lacking.

In this study, we wish to address this gap and have therefore implemented and assessed a large array of interpolation methods, aiming to better understand how different methods perform and which factors impact interpolation accuracy. To this end, we have collected several longitudinal datasets, each describing the composition of the gut microbiome of multiple individuals with different characteristics, and then used these datasets to assess each interpolation method. We specifically investigated how sample size, time intervals between adjacent samples, and host- and taxa-specific microbiome stability affect interpolation accuracy in each method. Finally, based on our findings, we present a simple approach to predict the expected interpolation accuracy in a given time point. We believe that such an analysis can benefit the microbiome research community, offering valuable insights and best practice guidelines for interpolating samples in longitudinal microbiome datasets. This, in turn, would facilitate the integration of such techniques into future studies, enhancing our capacity to understand the temporal dynamics of the microbiome.

## Results

### Evaluating longitudinal interpolation approaches

To estimate how well different interpolation approaches may work and what are the various factors that may impact interpolation accuracy, we first collected three 16S rRNA sequencing datasets, each describing the composition of the gut microbiome of multiple individuals over time. The first dataset was obtained from a study by Caporaso *et al*.^1^ and followed two individuals, a healthy male and a healthy female, for a long period of time (15 months for the male and 6 for the female), with an average sampling interval of 1.12 days. This is one of the densest and most long-term microbiome datasets available. The second dataset, published by Poyet *et al*.^14^, includes data from healthy fecal microbiota transplant donors. Here we used only individuals with at least 50 available samples, for a total of 9 individuals and 956 time points along 7-18 months. The third dataset was published by Muinck & Trosvik^4^, and followed 12 infants during the first year of life on a near daily basis. We annotated each sample at genus-level resolution, a commonly used practice in 16S sequencing-based datasets. To focus on taxa with more reliable measured abundance, we considered for each individual only the 30 taxa with the highest average abundance across all time points, with the abundances of all other taxa summed and classified in the analysis as ‘*others’*. Indeed, most of these additional taxa exhibit a markedly low prevalence (i.e., completely absent from many samples), an average relative abundance of <0.5% across all datasets, and ∼3.5% when aggregated together into the ‘other’ group, thus having a relatively limited impact on our estimation of interpolation accuracy.

We then implemented a large set of interpolation methods previously used in various studies. These methods can be roughly partitioned into four groups. The first group includes naïve interpolation methods^13,14^, in which the median abundance, the average abundance, or the abundance at the previous time point are used as the predicted abundance in a missing sample. The second group includes methods based on the generalized Lotka-Volterra (gLV) equations, commonly used to infer microbial interaction matrices^18–21^. These methods vary in how the species interaction matrix is inferred. Here, specifically, we used linear regression (which we term gLV MSE), MLRR^18^, and LIMITS^20^ (see Methods). The third group comprises general purpose machine learning and mathematical methods, including weighted average between the two adjacent time points, cubic spline^4,12,22^, and KNN (where closest neighbors are determined by proximity in time) with Epanechnikov kernel (see Methods). Finally, the last group is based on machine learning and statistical methods designed specifically for temporal analysis. Only a few methods from this group have been previously applied for temporal microbiome data modeling^12,13,15,16,23–25^. Here, we specifically used dynamic Bayesian network, as implemented in McGeachie *et al*.^26^ (CGBayesNets), using either a sparse and dense configuration (see McGeachie *et al*.^16^), which we term sparse DBN and dense DBN, respectively. As a “null” interpolation method, we further used a very naïve approach that assigns all the 30 taxa and the ‘*others*’ category an equal abundance (termed ‘equal’ in this work).

To examine how well each method can predict the composition of the microbiome in time points in which the microbiome was not assayed, we used a simple leave-one-out cross-validation approach. Specifically, for each individual in our datasets, we systematically omitted each available sample (except for samples taken at the first and last time points), and used each of the interpolation methods described above to infer the composition of the omitted sample based on all remaining samples. We then evaluated the degree of similarity between the inferred and real composition using the Bray-Curtis similarity metric to quantify the interpolation accuracy of each method.

### Variation in interpolation accuracy across datasets, individuals, time, and taxa

Following the methodology described above, we first examined whether certain methods generally perform better than others across all data or in certain datasets. This analysis suggested that KNN resulted in the best interpolation accuracy in all datasets, with mean similarity scores of 0.875, 0.833, and 0.77 in Caporaso et al., Poyet et al., and Muinck & Trosvik, respectively, for a mean similarity score of 0.8 across all datasets combined (Figure 1). Several other methods similarly exhibited high interpolation accuracy across all datasets; for example, weighted average showed good interpolation accuracy in all datasets, with a mean similarity score of 0.791 (second only to KNN). Both dense and sparse DBNs, as well as spline and MLRR, also exhibited relatively good accuracy overall. In contrast, both LIMITS and gLV MSE resulted in low similarity scores across all datasets, with mean similarity scores of 0.501 and 0.608, respectively. Specifically, interpolation based on gLV MSE resulted in the worst accuracy, second only to our null method, which assigned an equal abundance to all taxa.

**Figure 1:**
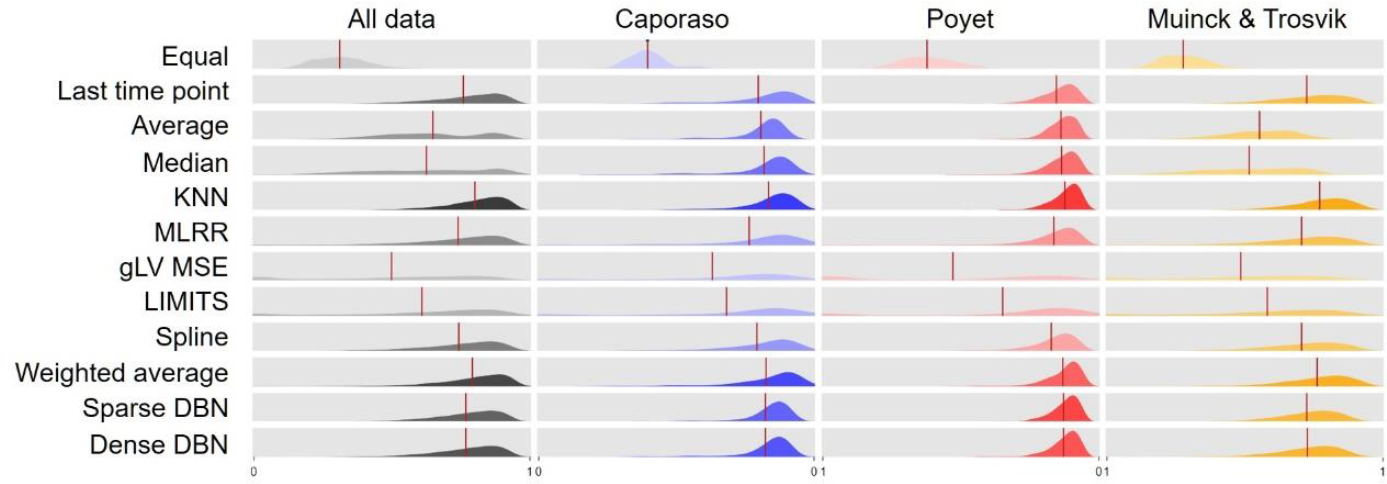
Interpolation accuracy across methods and dataset. Distribution of the Bray-Curtis similarity score between the interpolated composition and the real composition across all samples using a leave-one-out approach. Rows correspond to different interpolation methods and columns correspond to the different datasets (with the first column includes all the data, pooled). Red bars represent the mean value in each panel. The distribution’s transparency corresponds to the rank of the method’s mean within each dataset (i.e., as a measure of how accurate is each method in comparison to the others; more transparent means less accurate).

We also observed clear differences in interpolation accuracy between adults and infants cohorts within each method. Specifically, all methods, except for gLV MSE, performed substantially better in the data obtained from Caporaso et al. and Poyet et al. compared to the data obtained from Muinck & Trosvik (Figure 1), with a drop of 0.05-0.15 in the average similarity score in this dataset. This clear deterioration in interpolation accuracy in infants is perhaps not surprising given that microbiome composition during the first year of life tends to undergo significant changes^4,27^, making it harder to predict the composition of a missing sample based on the composition of samples in other time points. This deterioration in performance was especially noticeable in the naïve average and median methods that showed relatively good results in adults, with mean similarity > 0.8, but very poor results in infants, with mean similarity of 0.516 for the median abundance method and 0.553 for the average abundance method in the Muinck & Trosvik dataset.

Having established a clear difference in interpolation accuracy between adults and infants, we further examined whether a difference in accuracy exists also at the individual level. For this analysis, we have excluded the gLV MSE and LIMITS methods, which exhibited very poor interpolation accuracy above. Plotting the interpolation accuracy of each method for each individual clearly demonstrates variation across individuals (Figure 2). Moreover, as evident from this Figure, different methods vary similarly across individuals, with some individuals exhibiting relatively good accuracy across all methods, while others exhibit poor accuracy across all methods. Indeed, the correlation between the interpolation accuracy of the various methods across individuals was high (>0.9; pairwise Pearson correlation test). Moreover, this analysis further demonstrates that there is a clear and marked deterioration in performance in almost all infants and across all methods in comparison to adults, in agreement with the general decrease in interpolation accuracy in infants reported above.

**Figure 2:**
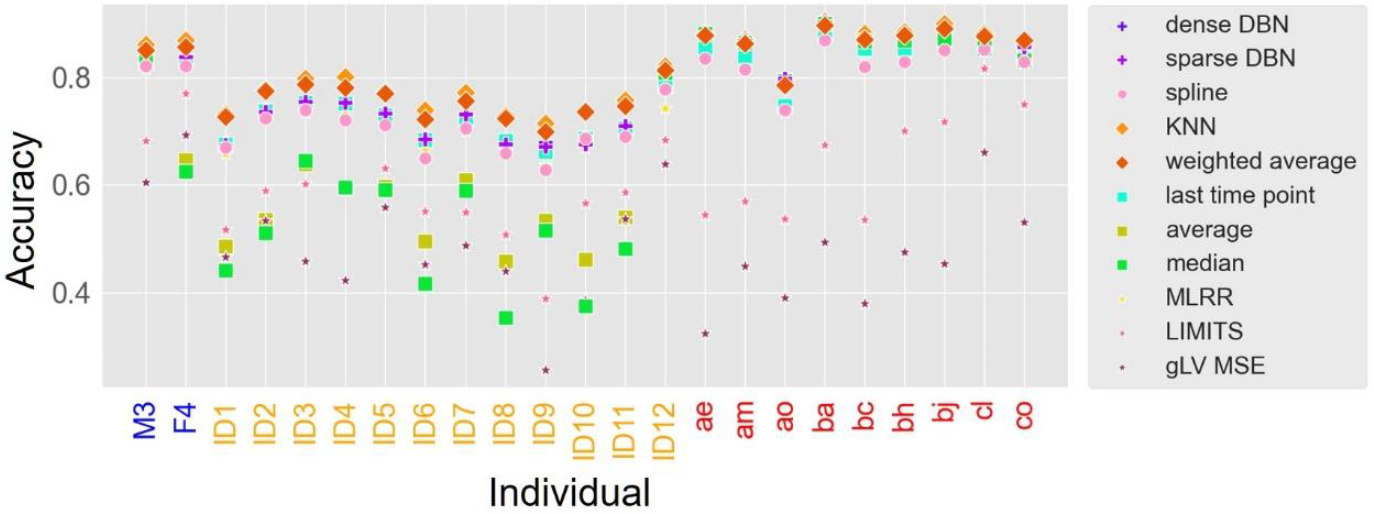
Interpolation accuracy vary across individuals. Average Bray-Curtis similarity between the interpolated and the real abundances, using a leave-one-out approach per individual. X-axis labels are colored by the dataset (red: Caporaso et al., blue: Poyet et al., orange: Muinck & Trosvik).

Finally, we set out to test whether interpolation accuracy also vary across different time point within the same individual. Indeed, examining interpolation performances in one example individual demonstrated such variation, again with different methods varying similarly across samples (Figure S1A). Moreover, examining the correlation between interpolation accuracy across samples obtained at time point of varying distances, suggested that interpolation accuracy is temporally auto-correlated. Put differently, the interpolation accuracy observed in a sample obtained at a given time point tends to be similar to the accuracy in samples taken at close time points, with this similarly decreasing with the distance between the two time points (Figure S1B).

Interestingly, beyond the variation in interpolation accuracy at the community level reported above, we also identified such variation across taxa within the community, wherein some taxa are easier (i.e., more accurate) to interpolate than others. To quantify the interpolation accuracy of a given taxon at a certain sample, we calculated the relative interpolation error for every taxa in each sample, i.e. 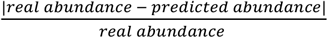. As depicted in Figure 3A, our analysis shows that, based on this metric, beside the naïve method (which assigns all taxa with equal abundance), the average, and the DBN based methods, all methods demonstrate similar behavior, both in terms of accuracy and stability in interpolation. Additionally, a noteworthy correlation can be observed between the average interpolation accuracy per individual at the community level (quantified above by Bray-Curtis similarity) and the average interpolation accuracy per individual at the taxa level (evaluated using the relative error; Figure 3B).

**Figure 3:**
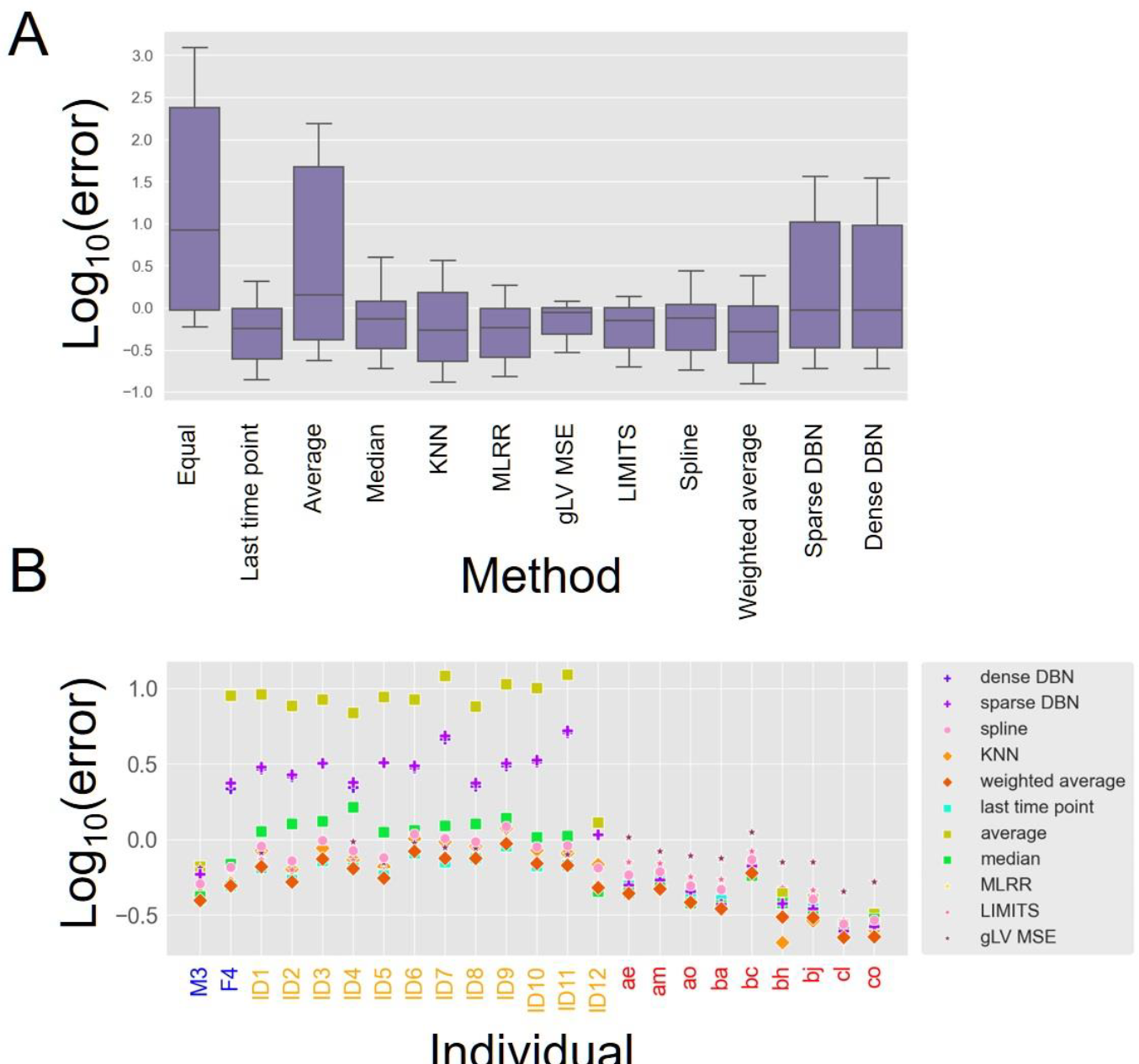
Relative error at the taxa level. **(A)** Distribution of the relative error across all taxa and in all individuals for each method. **(B)** Average relative error of all samples in each individual. Color of the X-axis labels represents the dataset (red: Caporaso et al., blue: Poyet et al., orange: Muinck & Trosvik).

Finally, following our observations above concerning the variability in interpolation accuracy across datasets, individuals, and time, we set out to more clearly compare and quantify the relationship between these axes of variation. As shown in Figure 4, there is indeed a strong correlation between the interpolation accuracy in a given time point and the interpolation accuracy in an adjacent time point from the same individual, and to a lesser extent, to the interpolation accuracy in some other time point from the same individual or in the same dataset. Interestingly, this finding suggests that it may be possible to *predict* the interpolation accuracy in a given time point using its neighbors’ inferred interpolation accuracy, as we indeed demonstrate later in this work.

**Figure 4:**
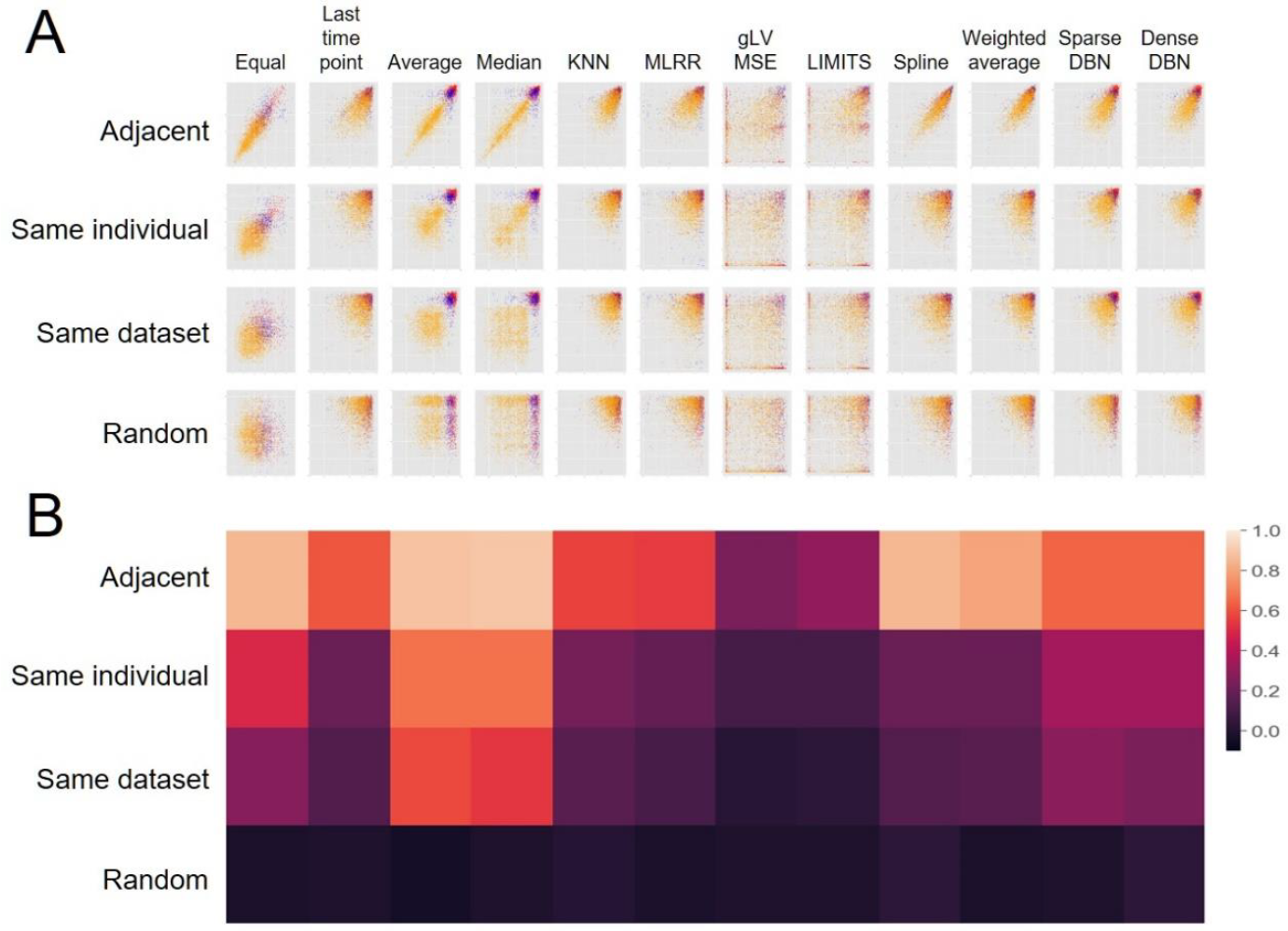
Correlation between interpolation accuracy within individuals, dataset, and adjacent time points. **(A)** The interpolation accuracy of each sample (using a leave-one-out approach; x-axis) as a function of the average interpolation accuracy in the adjacent time points (first row; y-axis), in a random time point from the same individual (second row), in a random sample from the same dataset (third row), and in a random sample from a random dataset (forth row). Columns represent the different interpolation methods and colors represent different datasets (blue: Caporaso et al., red: Poyet et al., orange: Muinck & Trosvik). **(B)** Pearson correlation coefficients of the scatter plots in Panel A.

### The impact of sample size and sampling frequency on interpolation accuracy

Given the marked variation in interpolation accuracy across datasets, individuals, and time demonstrated above, we next set out to examine whether specific properties of the data may additionally impact interpolation performances, focusing specifically on the number of available samples and on sampling frequency. Examining first the impact of sample size, we used a Monte-Carlo approach, randomly subsampling a certain number of samples from each individual (thus simulating individuals with varying sample sizes), and then randomly omitting one of these time points, interpolating the composition of this sample using the remaining samples in the subset of samples, and evaluating the interpolation accuracy as before. Repeating this process multiple times with different sample sizes allowed us to estimate the correlation between sample size and interpolation accuracy. We found, perhaps not surprisingly, that almost all methods (apart for the naïve approaches) exhibited at least some improvement with an increase in sample size (Figure 5), especially when the sample size was small. Furthermore, the improvement with sample size was particularly strong in MLRR and spline. Interestingly, our analysis also suggested that when the sample size is very small, simple weighted average method may outperform KNN.

**Figure 5:**
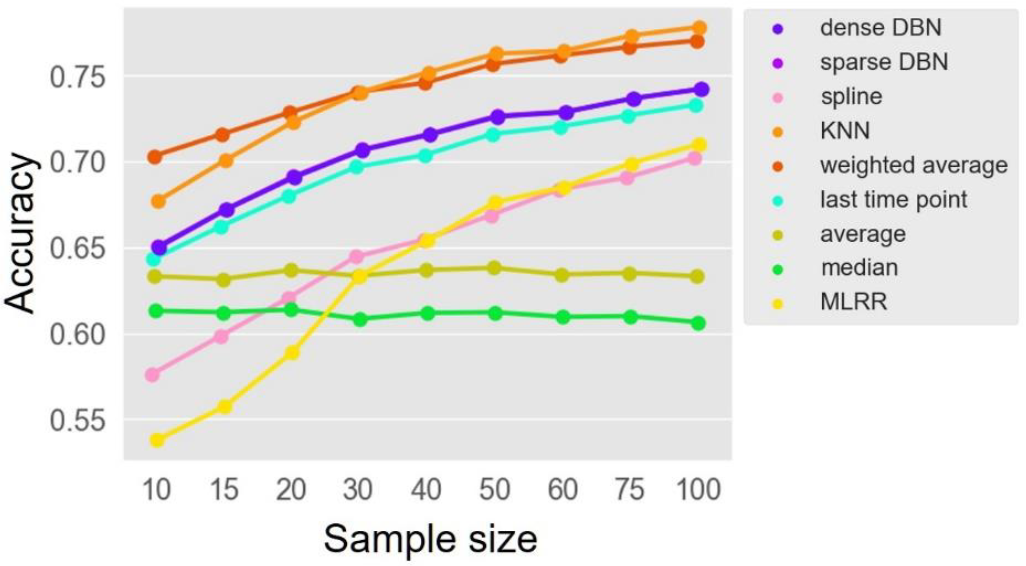
Correlation between sample size and interpolation accuracy. The effect of sample size on interpolation accuracy, measured using a Monte-Carlo approach.

Next, since our subsampling approach clearly induces a negative correlation between the sample size and the time interval between adjacent time points, we further examined whether the improvement in interpolation accuracy with sample size observed above is not purely due to the smaller interval between time points in the larger subsets. Specifically, to control for the possible confounding influence of one variable on the other, we fixed either the sample size or the time interval since the last sample, allowing only the other factor to vary. This analysis suggested that in most methods interpolation accuracy improves with both increased sample size and decreased time intervals (i.e., increased sampling frequency; Figure S2; Table S1). Some methods, however, did not follow these patterns, such as the naïve median and average, which is likely expected.

### Interpolation accuracy and microbiome stability

Clearly, our ability to accurately interpolate the composition of the microbiome at a given time point is tightly linked to the stability of the microbiome in that individual and around that time. Put differently, assuming microbiome stability changes across individuals and time^6,32,33^, when the microbiome becomes less stable, it is expected that interpolation accuracy will decrease at samples close to that time. Indeed, quantifying microbiome stability in each individual and around each time point (see Methods), we find it is significantly correlated with the interpolation accuracy calculated above (Figure S3). Interestingly the correlation between microbiome stability and interpolation accuracy was especially strong in the Muinck & Trosvik dataset, with most methods showing a spearman correlation > 0.5, potentially induced by the overall higher variation in microbiome stability in infants.

Next, we also wished to examine this relationship between interpolation accuracy and stability as the taxa level, hypothesizing that taxa with stable behavior, i.e. taxa whose relative abundance tends to stay constant over time, will be easier to interpolate. To this end, we used relative error as a measure for taxon interpolation accuracy, as described above, and similarly measured a taxon’s stability in a certain individual using the bimodality coefficient (as described in ^22^). High bimodality coefficient suggests that the taxon exhibits a non-stable behavior in a given individual (such as conditionally rare taxa or taxa with high fluctuations in abundance^22^). Indeed, comparing these two measures across taxa and individuals demonstrated a clear difference in the relative error between stable and non-stable taxa, with high bimodality taxa exhibiting higher relative error than low bimodality taxa (Figure S4).

Furthermore, we tested for a correlation between mean taxa stability in an individual and interpolation accuracy. To this end we compared the mean interpolation accuracy (measured using Bray-Curtis similarity score, as described above) of an individual and the mean bimodality of its’ most abundant microbiome taxa. Our analysis shows that in most methods, when the mean bimodality of the taxa in a given individual increases, the mean interpolation accuracy tends to decrease (P-value<0.01 in all methods except gLV MSE and <10^−7^in eight methods; Spearman correlation test).

### Predicting interpolation accuracy

Finally, we tried to harness our insights about interpolation accuracy reported above to construct an integrated interpolation accuracy predictor. To this end, we have fitted the data with a linear mixed effects model (LMM) with a random intercept, aiming to predict the logit product of the Bray-Curtis similarity score between the interpolation results and the real abundance. In our model, the random effect is the individual, and the fixed effect variables are the number of samples from that individual, the time difference from the preceding sample and to the subsequent sample, and logit transformation of the interpolation accuracy using Bray-Curtis similarity score in the preceding and subsequent samples.

P-value calculation for the fixed effect variables using Wald test showed that interpolation accuracy in adjacent samples is significant across all methods (Table S2). This is in agreement with our findings above concerning the autocorrelation in interpolation accuracy and the impact of time interval between adjacent time points (especially in methods that heavily rely on such temporal relationships, such as DBN, splines, and gLV). We also found that sample size was not significant in any method, and thus removed it from the model in downstream analyses. This in perhaps not surprising, since the sample size remains fixed within each individual and its impact is thus mitigated by the random effect in the model.

The model’s absolute error distribution further suggests that the linear mixed effect model described before is a reliable tool for predicting the interpolation accuracy, with mean absolute error below 0.1 across all methods (Table S2). Furthermore, Spearman correlation between the expected interpolation accuracy and the real interpolation accuracy was high (>0.6 in all methods), suggesting good performances of the model. Overall, these findings suggest that it is possible to predict the expected interpolation accuracy for a specific sample, offering researchers a way to not only interpolate the microbiome composition using the various methods, but also to assess the expected accuracy of this interpolation with high confidence.

## Discussion

This study compares and evaluates a series of interpolation methods for longitudinal microbiome data, aiming to identify the most suitable methods for predicting microbiome composition at time points when samples are missing. Such research, and the formulation of clear, best-practice guidelines for longitudinal microbiome imputations is essential given the recent rise in the prevalence of longitudinal microbiome studies. These studies often suffer from misaligned and incomplete sample collection, calling for effective interpolation of missing data that could facilitate more straightforward and standardized data comparison in downstream analyses. Furthermore, the unique characteristics of microbiome data, such as sparsity and high dimensionality, pose particular challenges for various interpolation methods, necessitating microbiome-specific method assessment.

Having implemented, explored, and evaluated a large array of interpolation methods, our results suggested that KNN with Epanechnikov kernel exhibits the highest interpolation accuracy in most settings. This observation is perhaps not surprising since compositional shifts in the microbiome are known to follow a gradual continuous process^5,6^, which Epanechnikov kernel KNN – an interpolation method that can be considered a generalized version of a weighted average – can successfully capture. As far as we know, however, this method has not yet been applied for longitudinal microbiome data interpolation, and we therefore encourage researchers to incorporate it in future longitudinal microbiome studies.

We also demonstrated that interpolation accuracy varies substantially across individuals, likely alongside variation in their microbiome stability (both at the individual level, and at the taxon level), with longitudinal data from some individuals supporting only very poor interpolation accuracy, regardless of the interpolation method used. This variation is especially marked in relation to age, as demonstrated by our finding that adults’ microbiomes can be more accurately interpolated compared to those of infants. Our analyses also revealed several other factors that affect interpolation accuracy, including sample size and sampling frequency.

Finally, demonstrating the practical impact our analysis may have, we have further used the insights we have obtained regarding interpolation accuracy to construct a model that successfully predicts, with high confidence, the expected interpolation accuracy at a given time point, based on linear mixed effect models. This model allows future studies to not only interpolate missing samples, but also to assess the expected interpolation accuracy in each time point and individual, and discard, for example, time points whose interpolated composition has low confidence from downstream analyses.

It is also important to note that the datasets we analyzed in this study include individuals whose microbiome was sampled for a long period of time and with a relatively high number of dense samples per individual. To date, unfortunately, most microbiome longitudinal studies are markedly less dense and include only a relatively limited number of samples per individuals. In this work we simulated such settings by subsampling our data. Despite this effort, however, there is likely still room for a similar comparative analysis focused specifically on smaller and less dense datasets. Furthermore, we only considered datasets that include healthy individuals and with no major perturbations over time. In such datasets, it is expected that the autoregressive regime of the microbiome will dominate its composition, an assumption that may not hold in datasets that include people with certain diseases or under certain medical or lifestyle interventions. Future studies could expand our analysis, including datasets obtained from individuals with various health conditions and examine interpolation methods that consider non-autoregressive perturbations.

More generally, we believe that collection of longitudinal microbiome data is of great importance to our understanding of the human gut microbiome, its dynamics, and its role in human health. Such data require carefully tailored computational methods for their analysis and research. We hope that the improved understanding of microbiome data interpolation gained in our work could be applied to such future studies and improve downstream analyses.

## Methods

### Interpolation methods

We have implemented and used an array of different interpolation methods (representing the four distinctive groups mentioned in the main text). These methods are described below.

First, we have implemented the naïve mean and median methods. These methods interpolate the abundance of each taxon using the mean or median abundance of that taxon across all available samples from the same individual. Another naïve interpolation method we have implemented simply uses the composition of the microbiome at the sample preceding the interpolated sample as the predicted composition at that sample.

An intuitive generalization of the last sample approach described above considers both the preceding and the subsequent samples, and uses a weighted average of these two sample with weights being inversed in proportion to the time interval between the interpolated sample and the preceding/subsequent samples. Formally, for a missing sample at time *t*, denote the closest preceding sample *x*_1_ at time *t*_1_, and the closest subsequent sample *x*_2_ at time *t*_2_. The predicted abundance at time *t* is then calculated as:

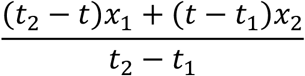

A more sophisticated generalization of this weighted average approach is based on the K-nearest neighbors (KNN) algorithm (nearest neighbors in time). This machine learning algorithm is often use to interpolate missing data in time-series anaysis^28,29^. KNN interpolates the microbiome’s missing composition *x* at time t using its *K* “closest neighbors”, i.e., the *K* known compositions, *x*_1_, …, *x*_*K*_, with times *t*_1_, …, *t*_*K*_ that minimize |*t* − *t*_*i*_| (in this study, we used *K =* 5). To consider the time differences of the samples used for interpolation from *t*, we weighted these samples based on Epanechnikov kernel. Overall, the algorithm calculates the composition *x* at time *t* in the following manner:

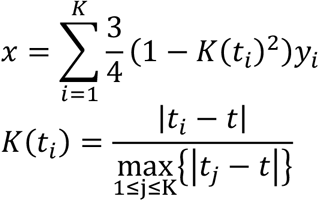

The resulting composition is then normalized to get a relative abundance profile.

Another mathematical interpolation method we considered in this work is based on a spline function. In this method, the abundance of each taxon at a missing sample is interpolated by fitting a cubic spline to its known abundance in all the other samples and using the obtained spline value at the time point with unknown composition to interpolate the taxon’s abundance. If the function output was negative for a certain taxon, the predicted abundance was set to 0. The resulting composition was then normalized as described in the KNN section. spline was calculated using Python’s Scipy^30^.

Dynamic Bayesian networks, a time-series specific machine learning model, could also be used for interpolation. For our study, DBN was represented as a bipartite directed graph. Specifically, for a dataset with n taxa, a set of n+1 nodes were generated, denoted here as 𝒜. The first *n* nodes of 𝒜 represent the community composition at a certain sample, *t* (i.e., each node denotes the relative abundance of one taxon) and the last node denotes the time difference between this sample and the subsequent sample. An additional set of *n* nodes, denoted here as ℬ, represent the community composition at the subsequent sample. All the edges in the graph are of form: {(*u, υ*)|*u* ∈ 𝒜, 𝒱 ∈ ℬ}, where edge (*u, υ*) represents the influence of taxon *u* at time *t* on the abundance of taxon 𝒱 in the subsequent sample. DBN structure and parameters inference was based on all pairs of non-missing subsequent samples using CGBayesNets^26^, and then applied to predict the missing composition based on the abundance in the previous sample. We have used both dense and sparse configurations as suggested in McGeachie *et al*.^16^ (these configurations are called here “sparse DBN” and “dense DBN” respectively).

Interpolation by ecological modeling was done using the generalized Lotka-Volterra (gLV) model. the gLV equations can be discretized as follow:

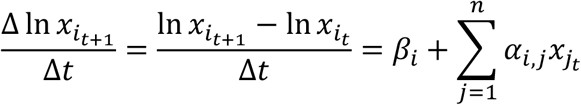

This equation represents the change rate in the abundance of one taxon. As there are many taxa and multiple time points, we can generalize this annotation, defining the matrix *F* as the matrix that represents this rate of change in abundance for all taxa in every time point simultaneously:

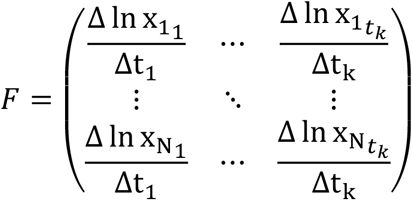

Then as demonstrated in Stein et al.^22^:

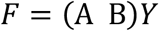

Where A_*i,j*_ *= α*_*i,j*_, B_*i*_ *= β*_*i*_ and

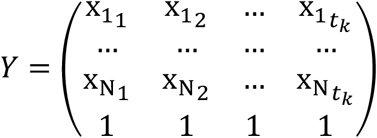

Since the matrix *Y* is known and since this is essentially the form of a regression problem, there are multiple ways to infer A and B. Here we used a simple estimator that minimized the MSE; maximum likelihood unconstrained ridge regression algorithm (MLRR) suggested by Stein et al.^18^ and LIMITS, suggested by Fisher et al.^20^. We have used the default parameters presented in the original papers.

It is worth noting that inference using the generalized Lotka-Volterra equations assumes that the time differences are all equal. Due to the characteristics of microbiome studies, this assumption does not hold in the datasets considered. Therefore, we have normalized *F* by dividing in the time difference between adjacent time points.

Finally, as a null reference, we have also implemented a naïve interpolation method that predicts all taxa to have the same abundance (called ‘equal’ throughout this work), meaning that for every taxon and at all time-points the interpolated abundance would be 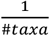

### Datasets and preprocessing

We utilized three longitudinal microbiome datasets with different characteristics that span common variability found across longitudinal microbiome studies.

The first dataset was obtained from a study by Caporaso *et al*.^1^ that followed two individuals, one healthy male and one healthy female, for a long period of time (15 months for the male and 6 for the female), with an average interval between samples of 1.12 days. The second dataset, published by Poyet *et al*.^14^, includes data from healthy fecal microbiota transplant donors. Here we used only individuals with at least 50 available samples from the dataset, for a total of 9 individuals and 956 time points along 7-18 months. Finally, the third dataset we used was published by Muinck & Trosvik^4^ and followed 12 infants during the first year of life on a near daily basis for a total of 2684 samples. As notes above, the microbiome undergoes significant changes during the first year of life, posing a particular challenge for any interpolation approach.

Feature tables for Caporaso *et al*. were obtained directly from the publication. Feature tables for Poyet *et al*. and Muick & Trosvik were calculated based on 16S read data published by the original papers. Specifically, processing of these data was done via the QIIME2^31^ pipeline, denoising was done using dada2^32^, the phylogenetic tree was constructed using q2-fragment-insertion^33^ plugin and SEPP^34^ technique, and taxonomy classification was extracted by q2-feature-classifier plugin^35^ based on a naïve-Bayes classifier and SILVA 99IU database^36^.

For each participant we considered only the 30 most abundant taxa across all time points, measured by the sum of the taxa’s relative abundance across all time points. The abundance of all taxa that were not among these 30 most abundant taxa, was summed and classified in our analyses as ‘*Others’*, as done in previous works^18,37–39^.

### Quantifying microbiome stability

Microbiome stability at a given time point was measured by averaging the Bray-Curtis similarity score between the microbiome composition at the two preceding time points, between the two successive time point and between the preceding and successive time points.

### Quantifying taxa stability

The stability of each taxon in every individual was measures using the *bimodality coefficient* as in Gibbons *et al*.^22^. Specifically, this bimodality coefficient, denoted as *β*, is calculated by the following formula: 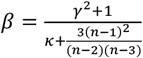, where *γ* is the empirical skewness, *κ* is the empirical kurtosis and n is the number of samples. Taxa exhibiting a *β* value greater than 0.7 were categorized as demonstrating stable behavior.

## Code availability

The Python code used for this analysis, including implementation of the various methods and analyses, is available on GitHub (https://github.com/borenstein-lab/longitudinal_interpolation).

## Acknowledgments

We would like to thank the Borenstein lab members for their helpful feedback and insightful discussions during the process. This work was supported in part Israel Science Foundation grant 2435/19 to EB. OP is supported in part by a fellowship from the Edmond J. Safra Center for Bioinformatics at Tel-Aviv University. The funders had no role in study design, data collection and analysis, decision to publish, or preparation of the manuscript.

## Supplementary Figures

**Figure S1:**
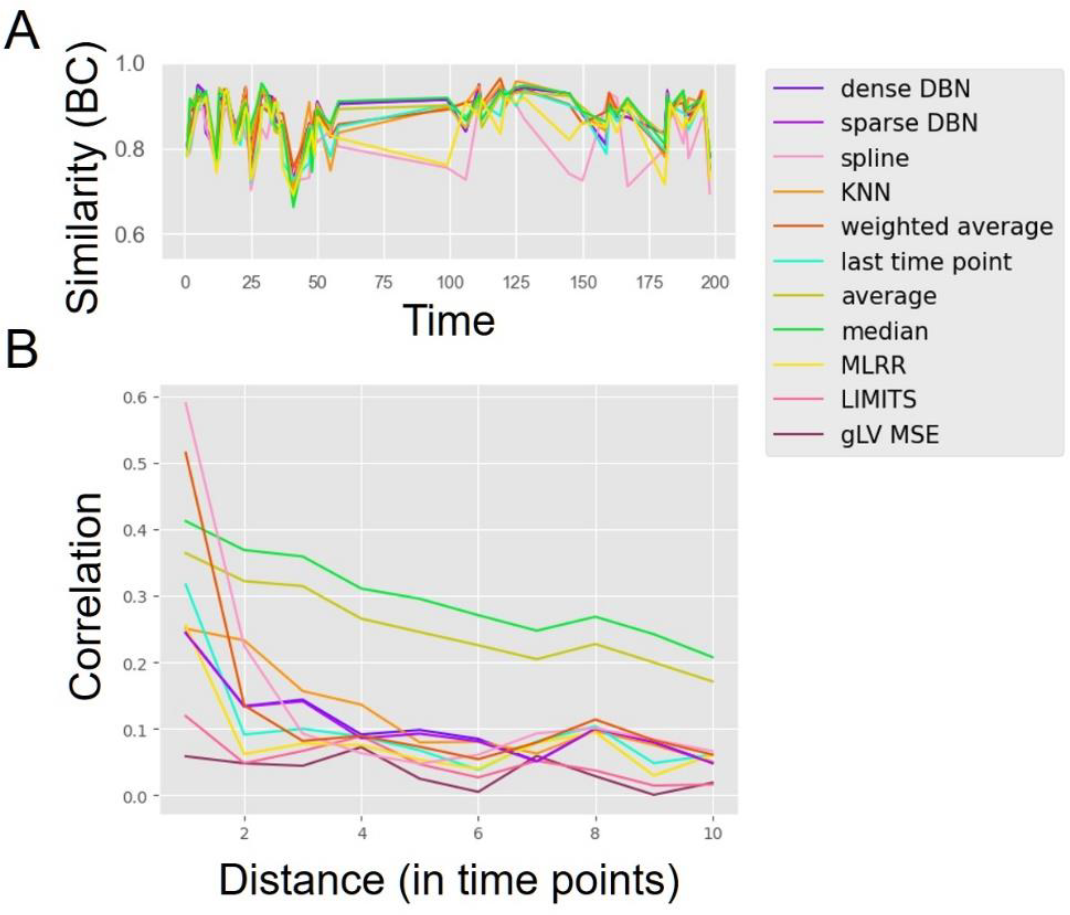
Interpolation accuracy across time. **(A)** Temporal interpolation accuracy for one specific individual (“ae”) from the Poyet *et al*. dataset. The lines represent the Bray-Curtis similarity between the interpolation results and the real abundance in each method. **(B)** The average auto-correlation over all individuals between interpolation accuracy of a sample and the interpolation accuracy of a preceding sample across varying time intervals.

**Figure S2:**
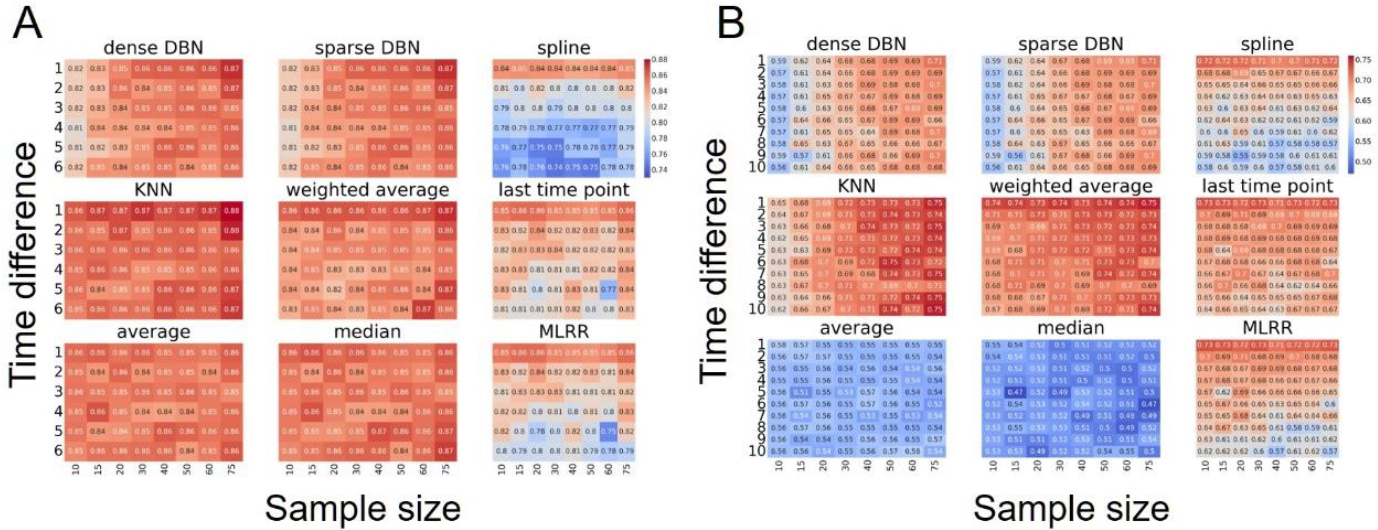
Interpolation accuracy is correlated with both sample size and the time interval between adjacent samples. Mean interpolation accuracy (across all individuals) for fixed sample sizes and time interval for **(A)** non-infants (Caporaso et al. and Poyet et al.2) and **(B)** infants (Trosvik & Muinck), considering only sample size and time difference pairs that had at least 50 samples.

**Figure S3:**
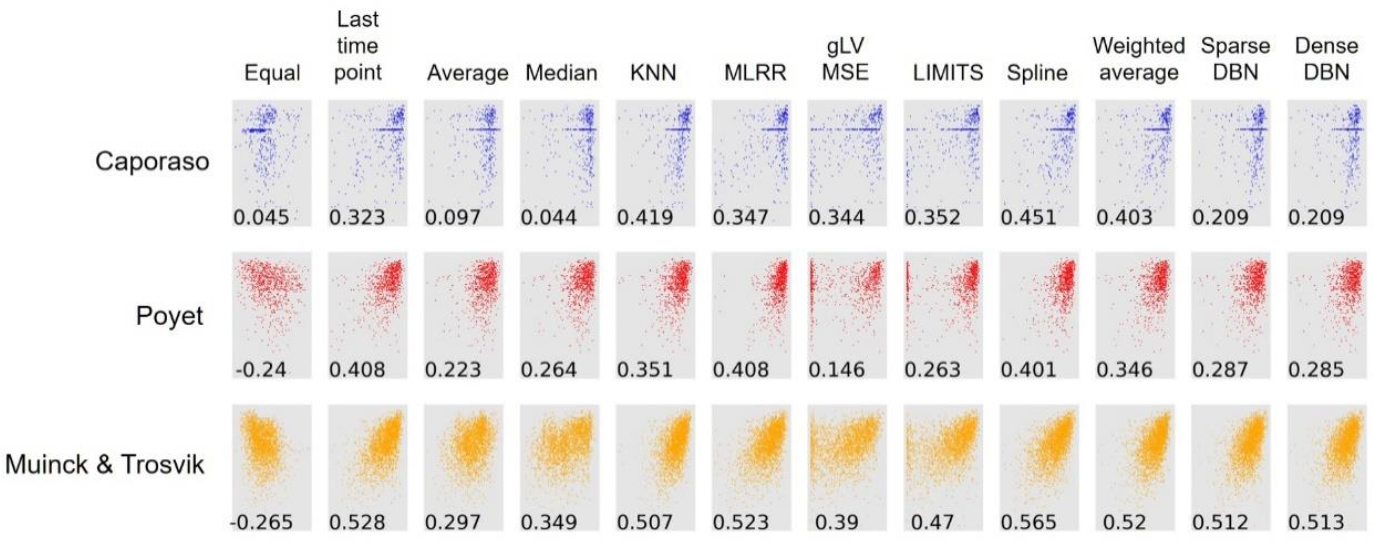
Interpolation accuracy and microbiome stability over time. Correlation between the interpolation accuracy at sample *i* and the stability of the microbiome around that sample (measured as the average BC distance between the samples at samples *i* − 2 and *i* − 1, *i* − 1 and *i* + 1, and *i* + 1 and *i* + 2). The Number at the bottom left of each panel denotes the spearman correlation.

**Figure S4:**
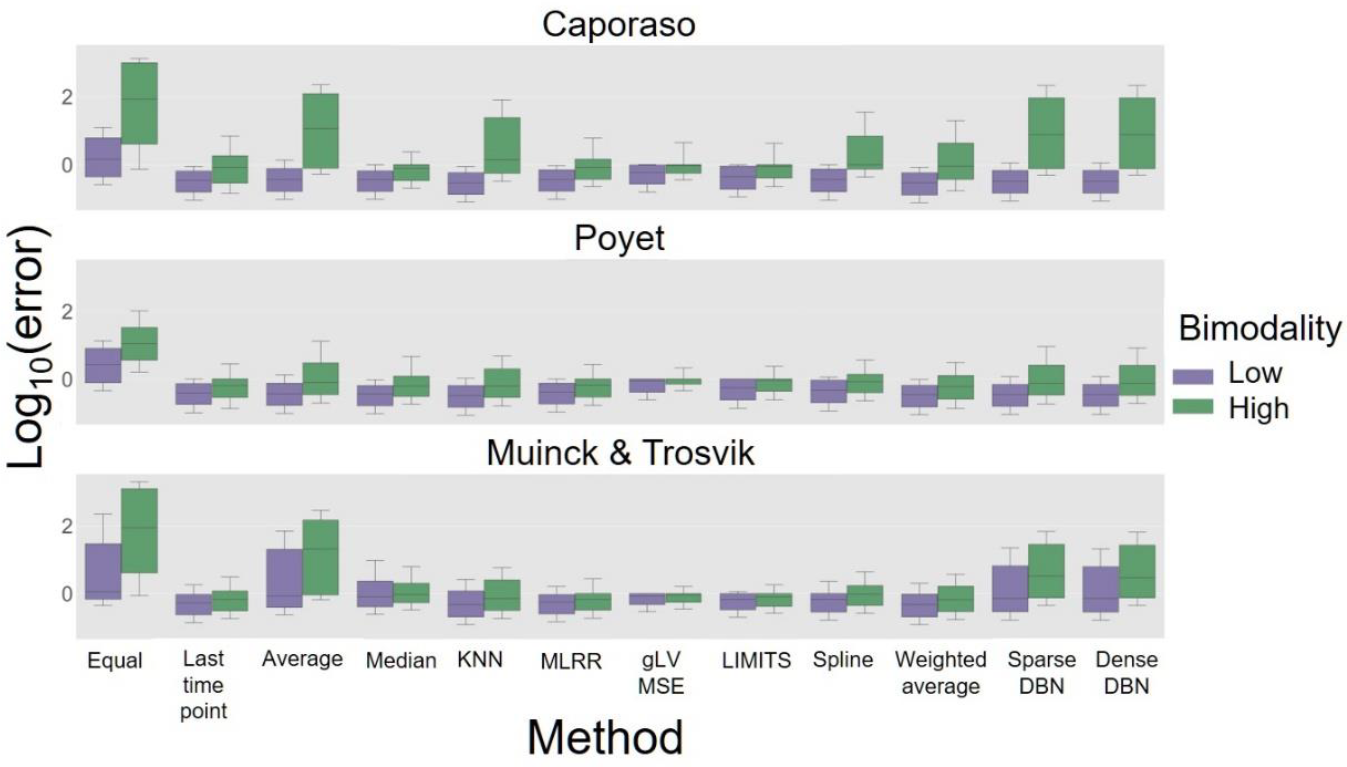
Taxa interpolation accuracy is linked to the stability of the taxa. Shown are boxplots denoting the distribution of relative error in taxa with low vs. high bimodality coefficient, for every taxon across all individuals and datasets. The cutoff for bimodality coefficient was set at 0.7, similarly to the 0.8 cutoff used in Ref^22^.

## Supplementary Tables

**Table S1:**
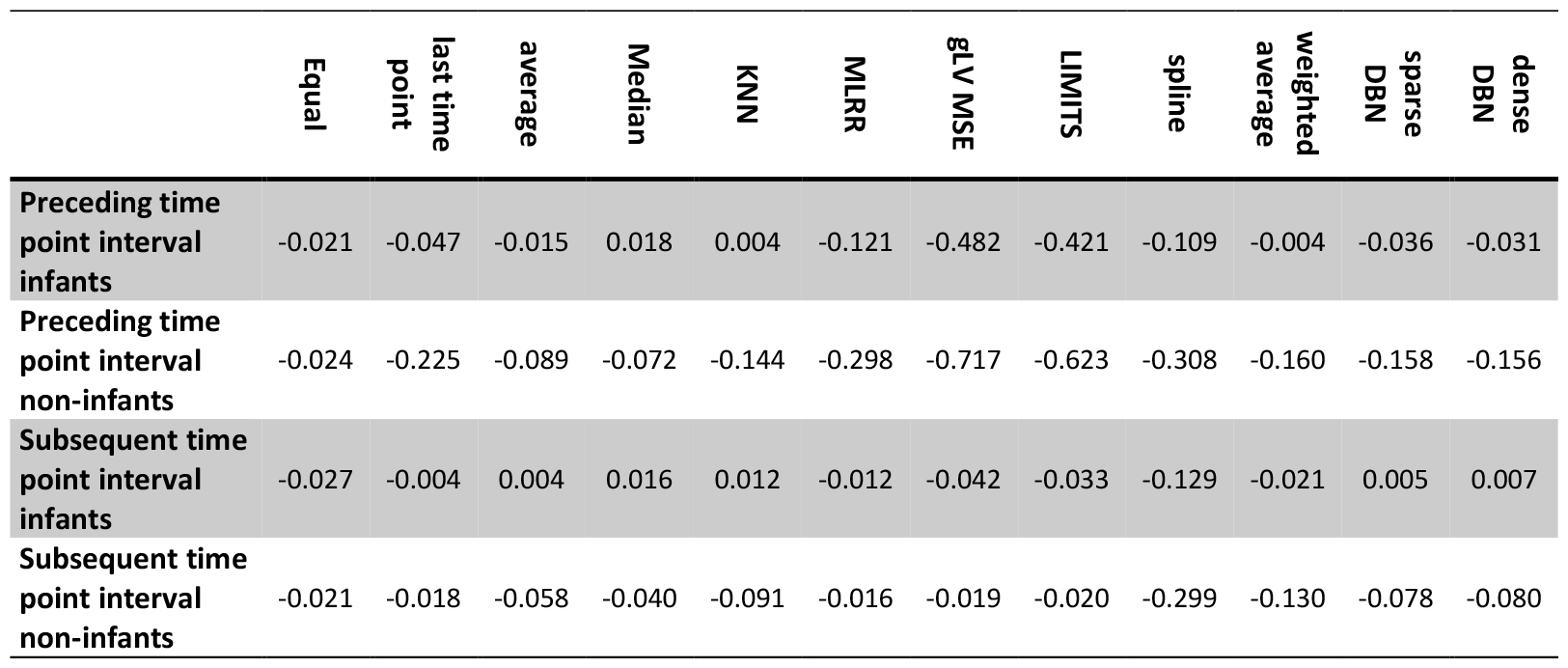
Spearman correlation between each method’s interpolation accuracy and the time difference from the preceding time point and the subsequent time point. Results are divided to infants (Muinck & Trosvik) and non-infants (Caporaso et al. and Poyet et al.).

**Table S2:** P-value for each variable in every method using Wald test for LMM model, as well as the mean and standard deviation of the absolute error between the model’s expected interpolation accuracy and the real interpolation accuracy across all time points.

|  | Equal | last time<br>point | average | Median | KNN | MLRR | gLV MSE | LIMITS | spline | weighted<br>average | sparse<br>DBN | dense<br>DBN |
| --- | --- | --- | --- | --- | --- | --- | --- | --- | --- | --- | --- | --- |
| **Preceding time**<br>**point time**<br>**difference** | 0.3374 | 1E-09 | 0.9019 | 0.3658 | 0.0263 | 3E-84 | 9E-159 | 1E-219 | 1E-12 | 0.1892 | 4E-05 | 5E-05 |
| **Preceding time**<br>**point quality of**<br>**interpolation** | 9E-223 | 4E-134 | 5E-140 | 6E-172 | 3E-91 | 2E-98 | 9E-06 | 2E-11 | 0 | 0 | 3E-99 | 2E-98 |
| **Samples count** | 0.6787 | 0.9008 | 0.3592 | 0.1993 | 0.6111 | 0.2304 | 3E-06 | 0.4575 | 0.0766 | 0.5419 | 0.5232 | 0.6009 |
| **Subsequent time**<br>**point time**<br>**difference** | 0.3031 | 0.4037 | 0.4604 | 0.5001 | 0.2955 | 2.3E-06 | 0.0035 | 0.3597 | 1E-22 | 0.0019 | 0.5556 | 0.6349 |
| **Subsequent time**<br>**point quality of**<br>**interpolation** | 2E-222 | 2E-134 | 5E-135 | 4E-167 | 3E-90 | 2E-103 | 9E-07 | 3E-14 | 0 | 0 | 3E-100 | 1.8E-99 |
| **Mean error** | 0.0370 | 0.0868 | 0.0730 | 0.0743 | 0.0667 | 0.0984 | 0.2242 | 0.1816 | 0.0530 | 0.0541 | 0.0709 | 0.0712 |
| **Error STD** | 0.0328 | 0.0831 | 0.0654 | 0.0735 | 0.0689 | 0.0992 | 0.1381 | 0.1378 | 0.0559 | 0.0527 | 0.0722 | 0.0720 |

